# Genomic Diversity, Host Associations, and Tissue Tropism of Hydrangea Ringspot Virus: A Global and Regional Perspective

**DOI:** 10.64898/2026.04.25.720767

**Authors:** Juliana Osorio-Marulanda, Juliana Lopez-Jimenez, Juan F. Alzate

## Abstract

Hydrangea ringspot virus (HdRSV) is an emerging plant virus infecting ornamental hydrangea species worldwide, yet its genomic diversity and host associations remain poorly understood. To expand the available genomic resources and assess HdRSV variability, we screened 210 publicly available *Hydrangea spp.* transcriptomes from diverse tissues, complemented with four newly generated *H. macrophylla* transcriptomes from Colombia. Viral genomes were assembled from infected samples and analyzed to infer phylogenetic relationships, lineage distribution, and relative viral RNA abundance. Two well-supported phylogenetic lineages (HdRSV-L1 and HdRSV-L2) were recovered from both full-genome and replicase coding sequence (CDS) analyses. HdRSV was detected across all host tissues examined, with the highest median viral loads in roots, followed by stems and leaves. *H. macrophylla* harbored both viral lineages, while *H. serrata* was exclusively infected by HdRSV-L1. Cultivar-level analysis revealed marked differences in viral abundance, with lineages showing distinct tissue preferences but no co-infection patterns, except in the ‘Bailer’ cultivar. Comparative analysis of the replicase CDS identified a single lineage-defining nonsynonymous mutation (C1578T; Thr→Ile), fixed in 90% of HdRSV-L2 genomes, corresponding to a polar-to-nonpolar amino acid change potentially associated with structural adaptation. Together, these findings provide the most comprehensive overview to date of HdRSV genomic diversity, host and tissue distribution, and molecular variation, offering new insights into the evolution and epidemiology of this understudied plant virus.

## INTRODUCTION

Herbaceous ornamental plants are valued for their aesthetic qualities and represent an important component of the global horticultural industry. They are used for both decorative and practical uses throughout the world, for that reason, they are very popular and economically relevant, and the market is growing every year. The production of herbaceous ornamental plants constitutes an international industry characterized by the large-scale trade of diverse species, most of which are propagated vegetatively. However, this intensive production system also exposes plants to phytosanitary risks that can compromise quality and lead to significant economic losses (Groth-Helms and Zhang, 2024; Shafiq et al., 2024).

Among these phytosanitary threats, viruses stand out as some of the most economically significant plant pathogens (Jones and Naidu, 2019). Plant viruses and viroids are a significant threat to ornamental species by reducing plant health and aesthetic value, leading to further economic impacts (Shafiq et al., 2024). This risk is amplified by the fact that the ornamental plant trade now represents a multibillion-dollar global industry (Hinsley et al., 2025). The industry’s massive scale is underscored by its valuation at $70 billion in 2024, drawing from a market spanning over 50 countries (Wakefield, 2025), with the global market forecasted to grow from USD 60.73 billion to USD 104.66 billion by 2033 (Market Growth Reports, 2025). The increasing globalization of trade, together with climate change and rising anthropogenic pressures, has intensified agricultural transformations that favor destructive viral disease outbreaks (Jones and Naidu, 2019). This intensified international exchange of plant material heightens biosecurity risks and accelerates the spread of infectious agents. In response, virus indexing programs are deployed to identify and eliminate infected material. However, the extensive diversity of both host species and their associated viruses complicate comprehensive detection, a challenge compounded by vegetative propagation practices, which can systematically perpetuate and disseminate (Groth-Helms and Zhang, 2024; Mitrofanova et al., 2018; Shafiq et al., 2024).

Within this context, *Hydrangea* species are particularly relevant ornamental hosts affected by diverse pathogens. The genus is susceptible to a broad range of diseases caused by organisms including the fungi *(Golovinomyces orontii*, *Cercospora hydrangea*, *Botrytis cinerea*, *Pucciniastrum hydrangeae*, *Colletotrichum gloeosporioides*, *Corynespora cassiicola*, *Phoma* spp., *Myrothecium roridum*, and *Alternaria* spp.), the bacterium *Xanthomonas campestris*, and various other viruses such as Hydrangea Mosaic Virus (HdMV), Tomato Ringspot Virus (ToRSV), and Tomato Spotted Wilt Virus (TSWV) (Li et al., 2016). However, among the viral pathogens, Hydrangea Ringspot Virus (HdRSV) is the main target for pathogen elimination programs in hydrangea cultivars (Tóth et al., 2012). First identified in 1952 (Brierley and Smith, 1952), HdRSV has since been reported in hydrangea cultivars across North America (Allen et al., 1983; Chiko and Godkin, 1986), Brazil (Van Sebroeck Dória et al., 2011), Europe (Bertaccini et al. 2015; Gadiou et al. 2010; Hollings 1958; Hughes et al. 2005; Mertelik y Kloudova 2009; Parrella y Vovlas 2004; Tóth et al. 2012), New Zealand (Pearson et al., 2006) and Asia (Li et al., 2020; Song et al., 2016; A Yusa et al., 2016). The global distribution of the virus is not precisely documented, but is probably present wherever hydrangea is cultivated as a result of the international trade of horticultural plants (Hughes et al., 2005a).

Taxonomically, HdRSV belongs to the genus *Potexvirus*, family *Alphaflexiviridae*, order *Tymovirales*, class *Alsuviricetes*, and phylum *Kitrinoviricota*, according to the International Committee on Taxonomy of Viruses (ICTV). In 2023, taxonomic adjustments included renaming the specie Hydrangea ringspot virus to *Potexvirus hydrangea* (ICTV 2025).

In 2025, the ICTV Master Species List (ICTV_Master_Species_List_2024_MSL40.v2) recognizes 52 species within the genus *Potexvirus*. These species are characterized by relatively short flexuous virions (<700 nm), primarily infecting herbaceous hosts, and lacking known biological vectors (ICTV 2025). Virions of potexviruses contain a single linear molecule of positive-sense RNA (5.9–7.0 kb), comprising 6 % of the virion’s weight. The RNA is capped at the 5′-terminus with m⁷G and contains a polyadenylated tract at the 3′-terminus (Fauquet et al. 2005; ICTV 2025; Kreuze et al. 2020). Typically, *Potexvirus* genomes encode five open reading frames (ORFs), but HdRSV is an exception with six ORFs: ORF1 (5′-proximal) encodes the replication-associated protein (replicase, 156 kDa); the subsequent triple gene block (TGB) comprises three overlapping ORFs that produce proteins (26, 12, and 8 kDa) for cell-to-cell movement; ORF5 (3′-terminal) encodes the coat protein (24 kDa); and ORF6, which is contained within ORF5, encodes a virus-specific protein (16 kDa) (Batten et al., 2003; Brunt, 2006; Gadiou et al., 2010a; Hughes et al., 2005a).

HdRSV infection can affect different tissues. Hydrangeas infected with this virus are often symptomless (Brierley and Smith, 1952; Nagashima and Tojo, 2023). However, in some plants, the presence of the infection has been associated with a range of manifestations, which can include chlorotic vein clearing, yellowish or leaves can become distorted, crinkled, or rolled. In severe infections, plants can exhibit systemic symptoms such as stunting, dwarfing, and leaf discoloration. It is also documented the reduction in the number and normal development of florets per inflorescence (Bertaccini et al. 2015; Gadiou et al. 2010; Koenig 1973; Parrella y Vovlas 2004; Tóth et al. 2012; Weiler 1980).

Historically, HdRSV has been diagnosed by detecting antigens with the double-antibody enzyme-linked immunosorbent assay (DAS ELISA; Adams y Clark 1977) and, more recently, molecular techniques like RT-PCR (Tóth et al., 2012). The complete genome of HdRSV was first determined in 2005 using Sanger sequencing (Hughes et al., 2005a). However, in 2025, only 18 HdRSV sequences are available in the NCBI Virus database. Most of these were generated using Sanger technology, four are less than 5300 bases, and despite the advent of more accurate high-throughput methods like Next-Generation Sequencing (NGS), most available data relies on older techniques.

To address this gap, we generated HdRSV genomic data using NGS (RNA-seq) from a symptomatic *Hydrangea macrophylla* plant cultivated in the Andean region of Colombia. This work provides the first genome sequence of HdRSV from Colombia and likely constitutes its first report in the country.

Furthermore, we conducted a large-scale in silico screening of 210 publicly available *Hydrangea spp.* transcriptomes from the NCBI Sequence Read Archive (SRA). The presence of HdRSV was confirmed in several datasets, enabling a broader assessment of its distribution. Subsequent phylogenetic analyses provided evidence for two distinct viral lineages, a finding reported here for the first time. We also performed association analyses to explore potential links between these viral lineages and factors such as geographic origin, host, cultivar, and tissue type. Finally, a transcript-level expression analysis offered new insights into viral activity, advancing our understanding of the molecular and evolutionary dynamics of HdRSV.

## METHODS

### 2.1. Plant Material and Sampling

The sampling was conducted at a commercial production farm of *Hydrangea macrophylla* located in the “Oriente Antioqueño”, subregion of Antioquia, Colombia, at an approximate altitude of 2,500 meters above sea level (m.a.s.l.). The commercial crop is cultivated within greenhouse conditions to ensure consistent and high-quality year-round flowering. Within this cultivation system, one plant exhibiting foliar necrosis and floral malformation was identified. This plant was subsequently transported intact and under live conditions to the laboratory for further evaluation, including subsequent RNA extraction procedures.

### 2.2. RNA-seq experiments from the Colombian hydrangea plant

In pursuit of identifying the etiological agent responsible for the observed alterations in the plant mentioned above, we conducted a detailed transcriptome analysis to search for viral agents that might be replicating in the Colombian hydrangea plant.

The plant of interest was processed at the CNSG laboratory, Universidad de Antioquia, Medellín, Colombia. Tissue segments (specifically leaf, stem, bud, and root) measuring approximately 2 cm×2 cm were selected and immediately flash-frozen in liquid nitrogen. The frozen tissues were then finely ground to a powder using a mortar and pestle under liquid nitrogen. A 100 mg aliquot of the resulting tissue powder was weighed and homogenized in 1 ml of Trizol™ Reagent. Total RNA purification was subsequently performed following the manufacturer’s standard protocol (Trizol™ Reagent, Invitrogen).

The sequencing was performed using the Illumina NovaSeq 6000 platform, with library preparation carried out using Illumina TruSeq kits and an rRNA removal strategy. The sequencing process generated paired-end reads of 100 base pairs (bp) in length. To ensure high-quality data, read quality assessment and trimming were performed using Cutadapt v3.5, a widely used bioinformatics tool for removing low-quality bases and adapter sequences (Martin, 2011). For transcriptome assembly, the Trinity software (version 2.13.2) was used (Grabherr et al., 2011).

The resulting contigs were then analyzed to identify potential HdRSV (Hydrangea ringspot virus) genomes. To perform this search, the BLASTN tool (version 2.12.0+) was used (Altschul et al., 1990; Zhang et al., 2000). A custom database was generated using HdRSV genome sequences downloaded from GenBank. Subsequently, contigs exhibiting significant homology hits (or similarity hits) were extracted. This was followed by the manual extraction and curation of the HdRSV sequences. While full genomes were retained, partial sequences were preserved and designated for complementary or supplementary analyses.

### 2.3. Hydrangea RNA-seq data from the SRA-NCBI database

A comprehensive set of hydrangea RNA-seq data was retrieved from the NCBI Sequence Read Archive (SRA). The data collection was performed via a targeted search for *Hydrangea spp.* transcriptomic projects, including all publicly available datasets deposited in the SRA up to September 2025. Selection criteria included Illumina-based sequencing, paired-end libraries and RNA source type. Application of these filters yielded 210 RNA-seq accessions along with their related metadata. Raw reads were processed to remove adapters and low-quality sequences, then assembled *de novo* using Trinity (Grabherr et al., 2011). For viral detection, contigs were initially screened for viral sequences, but the definitive confirmation of Hydrangea ringspot virus was based on mapping the reads to a reference genome (NC_006943.1). This sensitive mapping approach allowed for the reliable detection of the virus even in cases where the *de novo* assembly failed to produce viral contigs.

### 2.4. Enrichment of the dataset with publicly available sequence genomes for HdRSV

To enrich the genomic dataset and increase the robustness of the downstream phylogenetic analysis, publicly available complete HdRSV genomes were compiled from the NCBI Nucleotide database. These sequences were cross-referenced with the *Hydrangea spp.* sequences assembled from the SRA data to ensure a non-redundant final dataset. The final curated dataset consisted of 13 isolates, including the established reference genome (isolate PD_109, accessions AY707100.1 and NC_006943.1) and 12 additional isolates.

### 2.5. Phylogenetic analysis

For phylogenetic reconstruction, only sequences containing all six complete open reading frames (ORFs) of the HdRSV genome were retained, while incomplete or truncated sequences were excluded from subsequent analyses. Phylogenetic analysis was conducted using MAFFT v7.526 for multiple sequence alignment and IQ-TREE v3.0.1 for maximum-likelihood tree inference (Katoh and Standley, 2013; Wong et al., 2025). The analysis incorporated a total of 99 unique genomic sequences, 29 partial genomic contigs containing complete ORFs of HdRSV from the SRA dataset, one sequence generated in this study, 13 HdRSV publicly available sequences from NCBI and 53 representative *Potexvirus* genomes covering all recognized ICTV species. Three Lolium latent virus sequences were used as the outgroup to root the phylogenetic tree. The optimal nucleotide substitution model (TVM+F+I+R6) was selected using ModelFinder (Kalyaanamoorthy et al., 2017). Branch support was evaluated with 5,000 ultrafast bootstrap replicates (Hoang et al., 2018).

A complementary phylogenetic analysis was conducted focusing exclusively on the replicase coding sequence (CDS) to determine the lineage affiliation of HdRSV sequences. Phylogenetic reconstruction was performed under codon-based evolutionary models. Viral sequences containing at least 70% of the replicase CDS (703 codons; n = 123) were included, enabling the incorporation of 117 incomplete yet informative sequences in the phylogenetic dataset, together with the 13 HdRSV reference genomes available in GenBank and one additional GenBank sequence with >70% replicase coverage. The alignment was carefully curated to ensure that all sequences began at the canonical start codon (ATG) when the 5′ end of the CDS was present and that the correct reading frame was maintained across the entire region. The resulting phylogenetic tree was visualized and edited using FigTree software (Rambaut, 2018).

### 2.6. Global distribution and prevalence analysis

A global dataset was constructed by integrating the metadata of the newly generated viral sequences from SRA, this study, and all publicly available NCBI genomes. Prevalence and infection rates were calculated by tallying positive and negative detections for each country, host species, tissue type, and viral lineage. All analyses of geographic distribution and infection prevalence were conducted in RStudio (Posit, 2024) using the tidyverse package ecosystem (Wickham et al., 2019). The co-distribution of the lineages was visualized on a world map generated with the ggplot2, maps, and pattern packages (Becker et al., 2003; Wickham, 2016).

### 2.7. Quantification of viral genome abundance

To estimate the relative abundance of viral transcripts, de novo transcriptomes assembled with Trinity were processed to reduce redundancy by selecting the longest isoform per gene cluster. Cleaned paired-end RNA-seq reads were pseudo-aligned to the corresponding sample-specific transcriptome using *kallisto* 0.44.0 (default parameters, 20 threads), enabling transcript-level quantification without full alignment. Transcript abundances were normalized and expressed as Transcripts Per Million (TPM), allowing comparison across samples. TPM values were subsequently used to support comparative analyses of viral genome abundance across tissues, cultivars, host species, and viral lineages.

### 2.8. HdRSV expression and statistical analysis

Transcript abundance of HdRSV was quantified and normalized for the samples with successful *de novo* assemblies using Transcripts Per Million (TPM) via Kallisto v0.44.0 (Bray et al., 2016). The normalized TPM values, serving as an estimate of relative viral load, were integrated with a comprehensive metadata table. This combined dataset facilitated a comparative analysis of expression patterns across biological variables like host species, cultivar, tissue type, geographic origin, and viral lineage. Missing values were annotated as “NA” to keep dataset integrity. All statistical analyses and visualizations were conducted in R (v4.4.3) using RStudio (Posit, 2024). As the Shapiro-Wilk test confirmed, the data were not normally distributed across biological categories; non-parametric methods were employed. The Kruskal-Wallis test was used to infer global statistical differences across groups for each biological category, all of which returned significant results (p < 0.01). Subsequently, Dunn’s post-hoc test was applied for pairwise comparisons within these categories, and Wilcoxon rank-sum tests with continuity correction were additionally used in selected two-group comparisons where applicable. Data processing and statistical summaries were done using the tidyverse package collection (*dplyr*, *tidyr*), while visualization was implemented with ggplot2 (Wickham, 2016; Wickham et al., 2019).

### 2.9. Detection of HdRSV signature mutations

To identify lineage-defining mutations in HdRSV, the coding sequence (CDS) of the replicase gene was analyzed across all genomes assigned to lineages L1 and L2. Sequence alignments were generated using MAFFT v7.526 (Katoh and Standley, 2013) and manually curated in AliView v1.30 (Larsson, 2014) to ensure codon alignment consistency. Single-nucleotide polymorphisms (SNPs) were inferred using Treetime (Sagulenko et al., 2018) to detect fixed substitutions characteristic of each lineage. Both synonymous and non-synonymous changes were evaluated based on their lineage specificity. Nucleotide positions were reported relative to the replicase CDS coordinates of the reference genome NC_006943.

## RESULTS

### 3.1. Description of the Colombian *Hydrangea* plant

To investigate the presence of Hydrangea ringspot virus (HdRSV) in Colombian crops, we conducted field sampling in *Hydrangea macrophylla* cultivars near the municipality of Rionegro, Antioquia, located in the Andean region of Colombia. We surveyed plants exhibiting symptoms resembling the morphological alterations previously described for *H. macrophylla* infected with this virus. One plant displayed clear morphological abnormalities. Foliar examination revealed irregularly shaped necrotic lesions ranging in color from brown to reddish-brown, with a characteristic ash-grey center and distinct concentric rings. In addition, the plant showed reduced floral development, with only flower buds present and no fully developed inflorescences observed. This symptomatic plant was selected for RNA extraction and RNA-seq analysis, sampling leaves, roots, stems, and floral buds (Fig 1).

**Figure 1.**
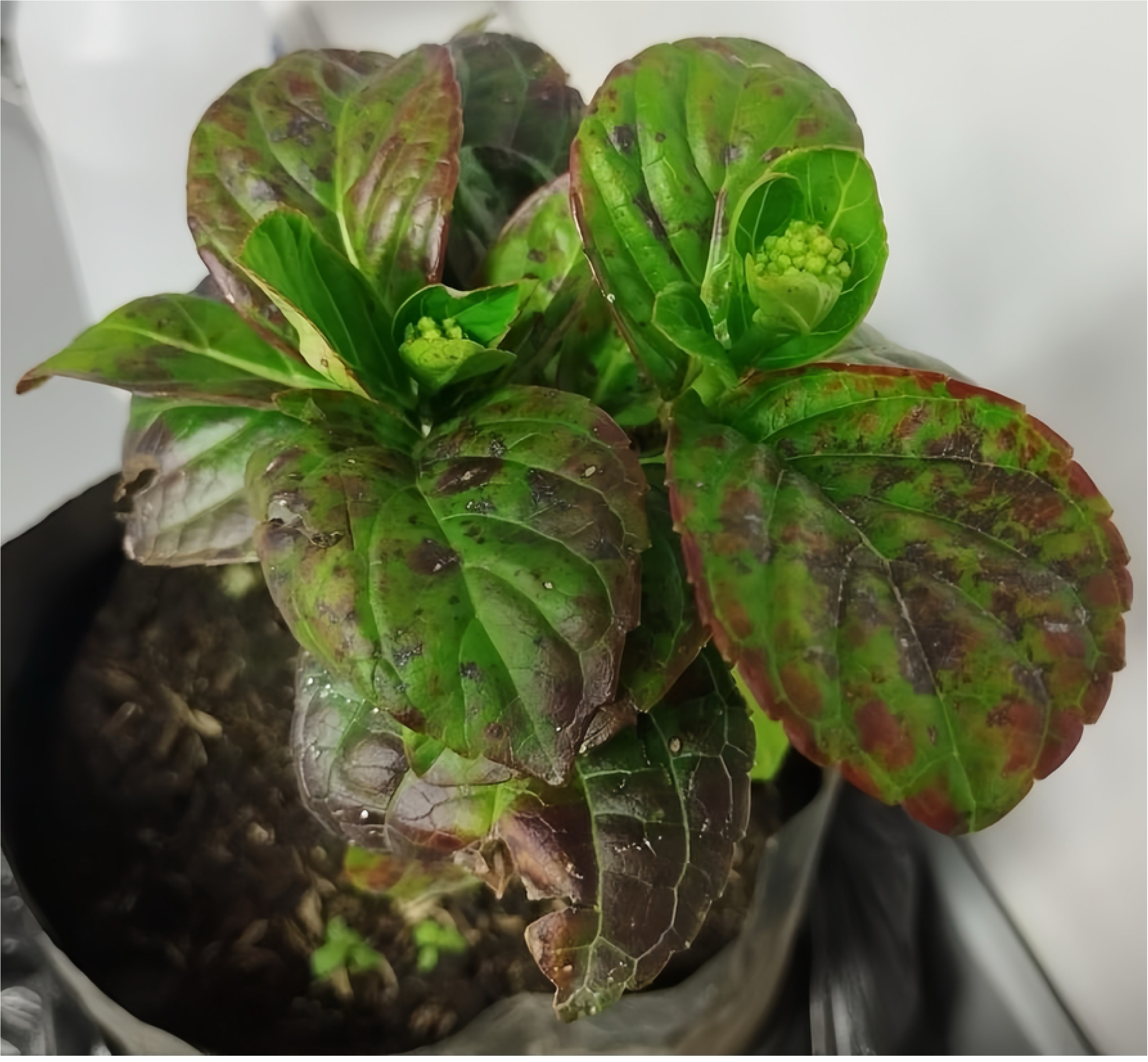
Morphological alterations observed in a Colombian *Hydrangea macrophylla*. The image shows foliar necrosis with concentric rings and ash-grey centers (yellow arrow), together with undeveloped floral buds (red arrow).

### 3.2. Generation of the first South-American HdRSV partial reference genome from a Colombian *Hydrangea macrophylla* plant

In order to confirm the presence of Hydrangea ringspot virus (HdRSV) in the Colombian *Hydrangea macrophylla* plant, a transcriptome analysis was performed using an RNA-seq strategy and *de novo* transcriptome assembly across four different tissues. Ribosomal RNA (rRNA)-depleted RNA-seq libraries were prepared from stem, leaf, root, and floral bud tissues. The root library yielded the largest dataset, comprising 6,162,932,130 raw bases and 61,019,130 paired-end reads. The bud library produced 5,323,617,484 raw bases and 52,709,084 paired-end reads, followed by the stem library with 5,248,335,114 raw bases and 51,963,714 paired-end reads, and the leaf library with 4,521,649,204 raw bases and 44,768,804 paired-end reads. Stringent quality control measures were applied to all raw sequencing data. Reads were filtered using a minimum Phred quality score threshold of Q30 and a minimum read length of 70 bases after trimming, ensuring that only high-quality reads were retained for subsequent analyses. The overall Q30 values were consistently high across all libraries, with 94.9% for stem, 94.2% for leaf, 94.5% for floral bud, and 95.3% for root.

*De novo* transcriptome assembly was conducted for the four tissue-specific datasets. The bud transcriptome assembly was the most extensive, yielding a total sequence length of 167.31 Mb, an average contig length of 723 bp, a largest contig of 25,278 bp, and the highest N50 value at 1,166 bp. The stem assembly followed in overall size, reporting a total length of 139.19 Mb, an average contig length of 595 bp, a largest contig of 20,285 bp, and an N50 of 849 bp. Conversely, the assemblies for root and leaf were shorter. The root assembly reached a total length of 82.21 Mb, an average contig length of 585 bp, a largest contig of 22,086 bp, and an N50 of 831 bp. Finally, the leaf assembly, while having the lowest total length (76.82 Mb), achieved a high degree of contig continuity, evidenced by the largest overall contig (32,375 bp), an average contig length of 665 bp, and an N50 of 1,046 bp.

The search for Hydrangea ringspot virus (HdRSV) sequences was performed using the BLASTN tool against a custom database containing GenBank reference genomes of the virus, as described in the Methods section. Within the *de novo* assembled transcriptomes of the different tissues from the Colombian *Hydrangea macrophylla* plant, three tissues—stem, root, and floral bud—yielded positive HdRSV contigs, whereas the leaf transcriptome showed no viral hits. The largest viral contig, measuring 4,331 base pairs (bp) (average sequencing depth 20.8X), was identified in the stem transcriptome and encompassed approximately 70% of the HdRSV genome, suggesting a higher viral load or more active replication in this tissue. In contrast, assemblies from the root and bud tissues produced substantially shorter viral fragments, measuring 2,343 bp (average sequencing depth 25.2X) and 632 bp (average sequencing depth 1.3X), respectively, none of which allowed the reconstruction of the complete viral genome. The absence of HdRSV contigs in the assembled leaf transcriptome and the partial assemblies in root and bud suggest tissue-specific variation in viral replication rate (Supplementary Table 1).

The primary contig retrieved from the stem transcriptome—the longest among all assemblies—was further analyzed and found to encompass the viral replicase coding sequence (CDS), specifically corresponding to Open Reading Frame 1 (ORF1). This contig showed a high nucleotide identity of 96% when aligned to the reference Hydrangea ringspot virus (HdRSV) genome (GenBank accession NC_006943.1). The viral contig was supported by 617 mapped reads, resulting in an average sequencing depth of 20.8×.

Subsequent manual annotation and curation of this transcriptomic contig confirmed a complete replicase CDS, including the correct start codon (ATG), appropriate coding sequence length, and the presence of the expected stop codon. These findings validate the integrity of the assembled viral sequence.

This study reports the first partial genomic sequence of Hydrangea ringspot virus (HdRSV) identified in South America. The 4,331-base pair (bp) partial genome, assembled from the stem transcriptome of a Colombian *Hydrangea macrophylla* plant, has been deposited in the NCBI database under the name HdRSV isolate UdeA (Accession number pending). As addition support of this finding, we also detected the presence of viral sequences (transcriptomic contigs) in root and floral bud transcriptomes of the same plant.

### 3.3. Identification of novel HdRSV genomic sequences from *Hydrangea* RNA-seq data

To expand the available genomic resources for HdRSV, we screened 210 publicly available Hydrangea transcriptomes derived from diverse tissues, complemented by four newly generated transcriptomes from the Colombian *H. macrophylla* plant. The resulting dataset comprised leaf (n = 66), stem (n = 28), floral structure (sepals and inflorescence; n = 39), bud (n = 37), and root (n = 43) transcriptomes. Publicly available raw RNA-seq data were retrieved from the NCBI Sequence Read Archive (SRA) and processed through quality filtering, adapter trimming, and de novo transcriptome assembly.

Using the BLASTN-based detection strategy, HdRSV-positive contigs were identified in 72% of the transcriptomes (155 samples), indicating a broad viral presence across samples. To increase detection sensitivity, we subsequently performed read mapping against the HdRSV reference genome (NC_006943.1), which revealed 166 positive RNA-seq samples (with at least 1 high-quality read mapped to the reference), eleven more, increasing the percentage of positive samples to 77.6% (Supplementary Table 1).

In several cases, the Trinity assembler produced two to five HdRSV genomic contigs per sample, typically covering different regions of the viral genome. For 19 samples, nearly full-length HdRSV genomes were successfully reconstructed, including the complete coding sequences (CDS) of all viral genes. These assemblies were only partially incomplete in the untranslated regions (UTRs), where small stretches of bases were missing. Interestingly, in samples with low sequencing depth (<20×), the assembled viral contigs were typically limited to regions encompassing the replicase CDS and other coding sequences located near the 3′ end of the viral genome. In contrast, it was often difficult to recover contigs spanning the genomic region between nucleotides 4,250 and 5,250 in this low coverage samples. To investigate whether a sequencing bias affected this region, we performed a coverage mapping analysis using the HdRSV reference genome (NC_006943.1). This analysis revealed a notably lower sequencing depth in that genomic interval across all positive samples. Furthermore, a GC content profile of the HdRSV genome, in windows of 200 bases, showed that this same region exhibited a prolonged peak of elevated GC content, suggesting that the reduced coverage may be related to sequence composition bias affecting library preparation or sequencing (Fig 2).

**Figure 2.**
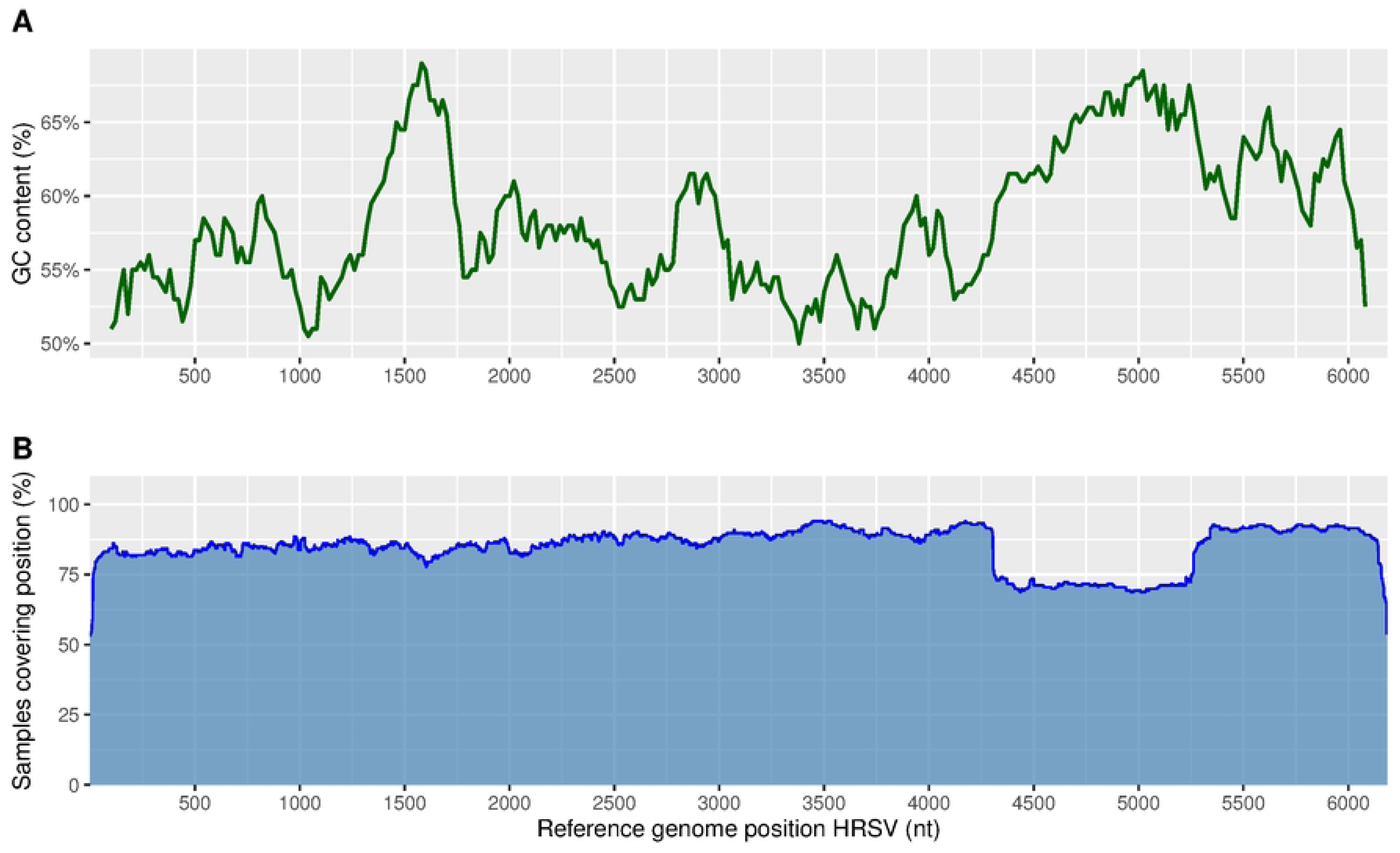
GC content and sequencing coverage profile across the Hydrangea ringspot virus (HdRSV) reference genome (NC_006943). (A) GC content (%) calculated in sliding windows of 200 bp with 20 bp step size along the HdRSV genome, showing a pronounced GC-rich regions. (B) Percentage of HdRSV-positive samples with sequencing coverage at each viral genome position, based on read mapping to the reference genome.

To validate the non-redundancy of our dataset, the SRA entries corresponding to the assembled viral genomes were cross-referenced with the HdRSV reference sequences available in GenBank. This comparison identified a linked GenBank record and its corresponding SRA RNA-seq dataset originating from a *Hydrangea macrophylla* sample collected in South Korea. Both entries were intentionally retained: the SRA-derived sequence was used for downstream expression and abundance analyses, while the GenBank reference genome was included in the phylogenetic reconstruction.

### 3.4. Phylogenetic analysis delineates two major HdRSV lineages

To gain insights into the evolutionary history of HdRSV, a phylogenomic reconstruction of the genus *Potexvirus* was conducted using Lolium latent virus as an outgroup. The analysis included reference genomes representing all recognized *Potexvirus* species (n = 53), together with 13 HdRSV genome references available in GenBank and the viral genomic sequences *de novo* assembled in this study, each containing at least the complete coding sequence (CDS) of the viral replicase gene (n = 30), including the newly characterized Colombian isolate designated HdRSV-UdeA.

The maximum-likelihood phylogenetic reconstruction revealed two well-supported evolutionary lineages, designated HdRSV lineage 1 (HdRSV-L1) and HdRSV lineage 2 (HdRSV-L2). The segregation of these clades was strongly supported, with ultrafast bootstrap values of 98 for HdRSV-L1 and 100 for HdRSV-L2 (Fig 3). A single sequence from a *Phlox* host collected in Germany did not cluster with either lineage and instead occupied the most basal position in the tree, suggesting it may represent an early-diverging HdRSV variant.

**Figure 3.**
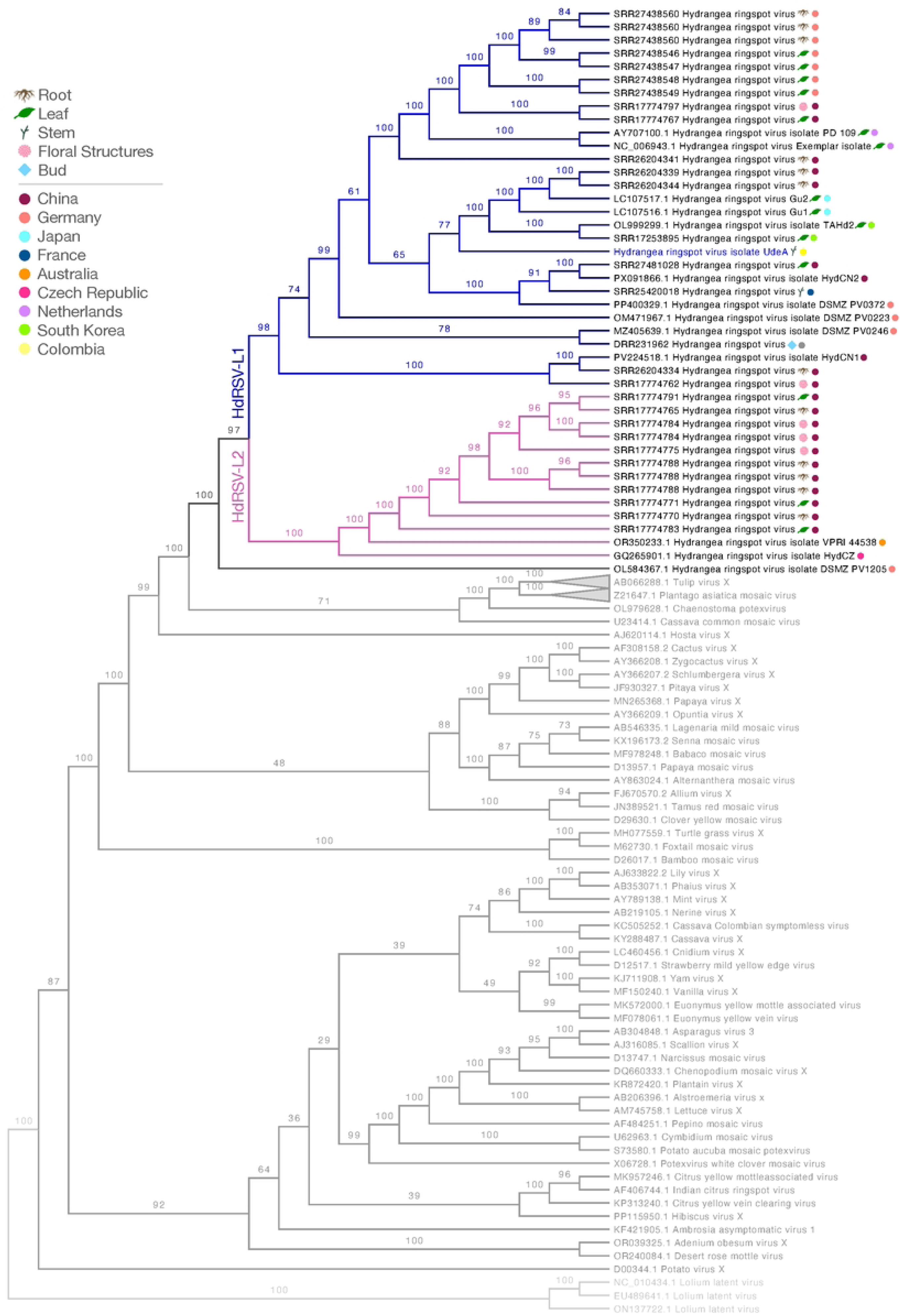
Maximum-likelihood phylogenetic tree based on sequences containing complete coding regions. The analysis includes HdRSV and other recognized species within the genus *Potexvirus*. Lolium latent virus was used as the outgroup for rooting. The scale bar represents transformed branch lengths, not genetic distances. Numbers at nodes indicate UFB support values from 5000 pseudoreplicates.

As a complementary approach, we conducted a second phylogenetic analysis focused exclusively on the replicase coding sequence (CDS), employing codon-based evolutionary models. This reconstruction included viral sequences from samples containing at least 70% of the replicase CDS (703 codons; n = 123), together with 14 HdRSV genomes available in GenBank (S1 Fig).

### 3.5. Global distribution and host range of HdRSV lineages

Using metadata available from the NCBI Sequence Read Archive (SRA), together with information from GenBank reference records, we analyzed the geographical distribution of HdRSV infections and the hydrangea species associated with viral presence based on RNA-seq data. Furthermore, we examined the global distribution of infections corresponding to the two phylogenetic lineages identified in this study, HdRSV-L1 and HdRSV-L2.

Geographical analysis revealed distinct yet partially overlapping distributions for the two lineages. HdRSV-L1 was detected in most analyzed regions but was absent from the only Australian dataset. In contrast, HdRSV-L2 was identified in Germany, China, and Australia. Both lineages co-occurred in Germany and China; notably, in China, HdRSV-L1 and HdRSV-L2 were detected in replicate samples of the same hydrangea cultivar, ‘Bailer’ (Fig. 4).

**Figure 4.**
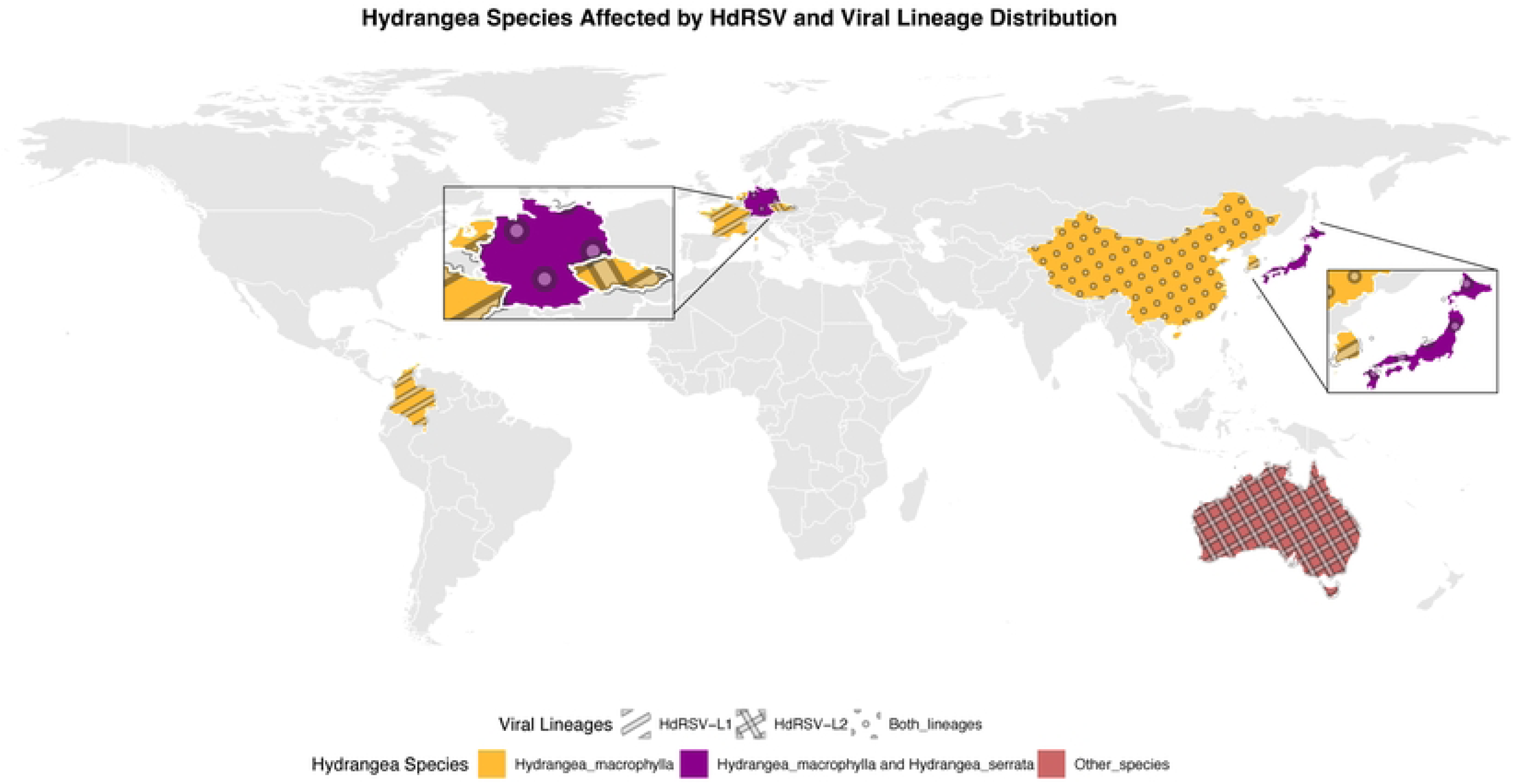
Geographic distribution of HdRSV genetic diversity. This world map integrates newly generated data from SRA analysis, a novel sequence from Colombia, and all publicly available NCBI genomes to show the global occurrence of HdRSV. The map displays the co-distribution of the two phylogenetic lineages (HdRSV-L1 and HdRSV-L2) and the host species from which each virus sequence was obtained.

The host species distribution showed that most HdRSV-positive sequences were from *H. macrophylla*, followed by *H. serrata* and other species (Fig 5). The overall infection rate was high (around 78%), although virus-free specimens were confirmed. The dataset was predominantly composed of samples from China, Germany, Japan, and France. These results reflect the availability of public transcriptomes rather than true regional prevalence.

**Figure 5.**
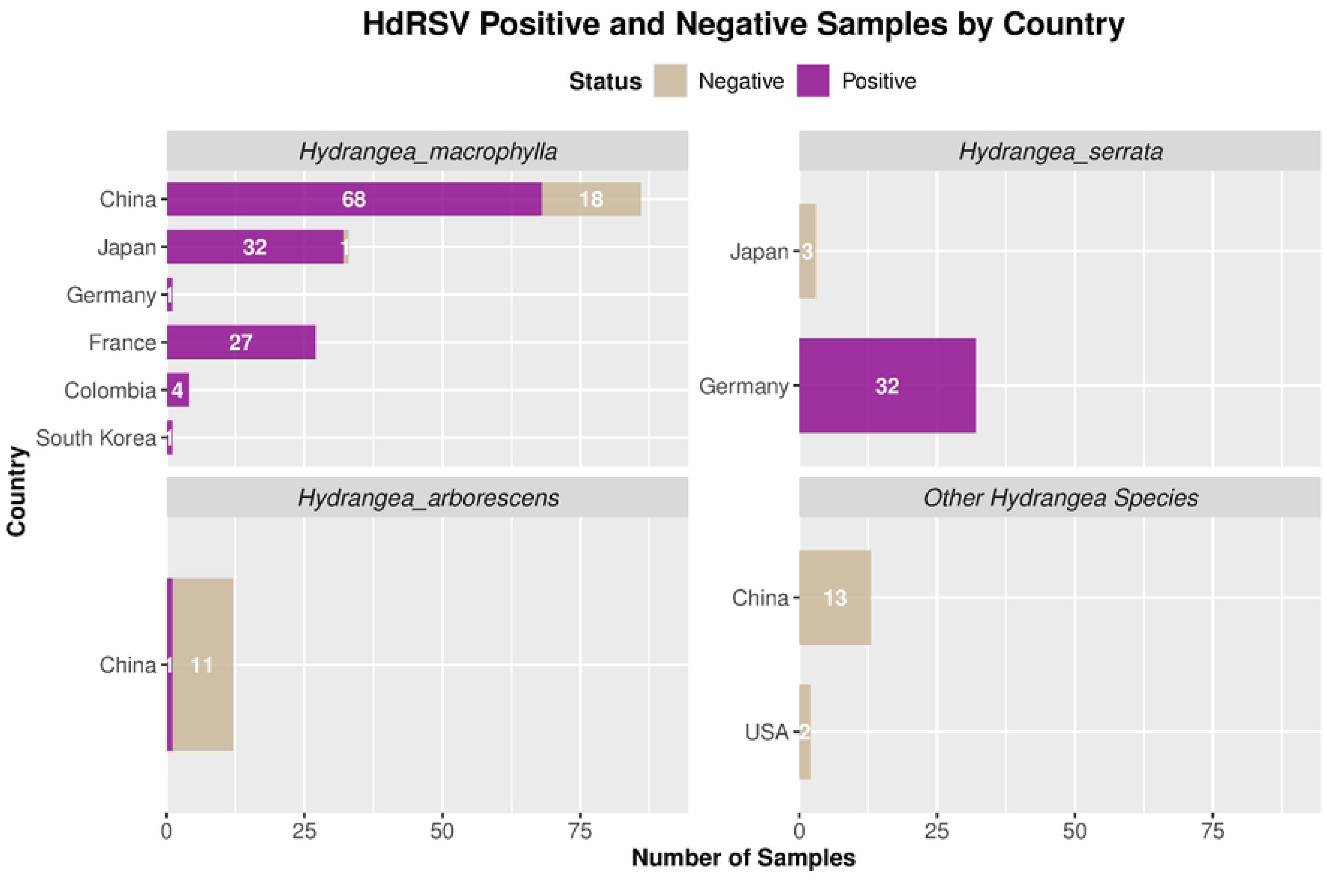
Prevalence of HdRSV in public SRA data and new Colombian samples. This bar chart summarizes the outcome of HdRSV screening, incorporating public Sequence Read Archive (SRA) datasets and new positive detections from Colombia. The results are grouped by country and host species, showing the proportion of positive and negative detections within each group.

### 3.6. Association of HdRSV with host tissue, cultivar, and lineage

HdRSV was detected in transcriptomes from all host tissues analyzed in this study. The highest number of HdRSV-positive assemblies was found in leaf transcriptomes (n = 66), which also represented the most abundant sample type in our dataset—reflecting the greater availability of leaf-derived RNA-seq data in the SRA database. Stems exhibited the highest positivity rate, with 100% of the samples (n = 28) testing positive for HdRSV, followed by roots, which were the second most represented tissue type and showed a positivity rate of 95.3%. In contrast, floral structures exhibited the lowest detection rate at 51.3%. *Hydrangea macrophylla* was the only species represented across all tissue types, whereas *H. serrata* was limited to leaf and root samples, both of which were predominantly infected (Fig 6).

**Figure 6.**
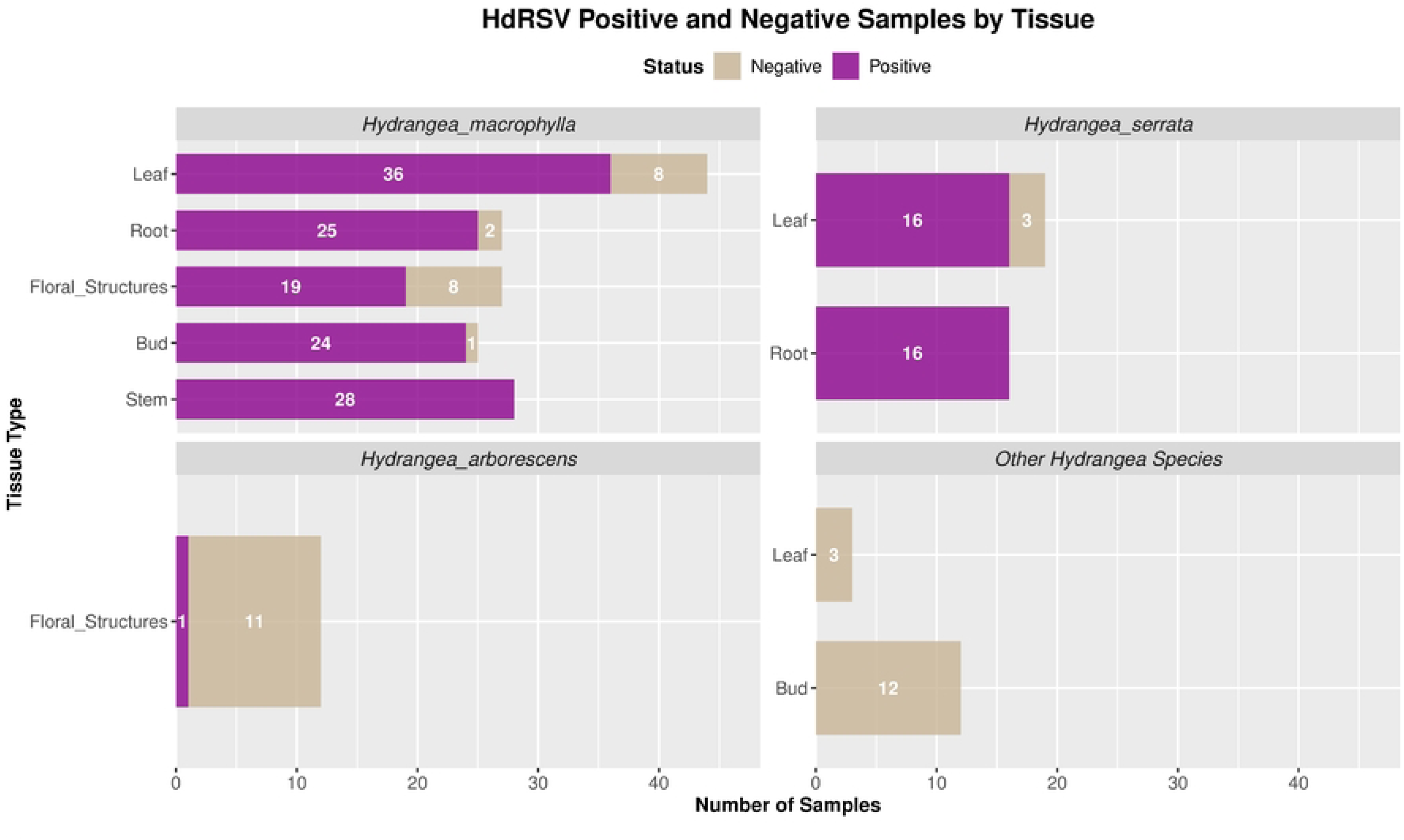
Tissue-specific prevalence of HdRSV. Based on data from SRA and this study (including data from Colombia), this chart shows virus detection results across different tissues, grouped by host species. Complete tissue representation (root, leaf, floral structures, and stem) is only shown for *H. macrophylla* due to data availability constraints for other species.

When comparing *Hydrangea* specimens segregating them by cultivar, plants of a given variety were typically uniformly infected (either all positive or all negative), with mixed infection status within a cultivar being less frequent (Fig 7).

**Figure 7.**
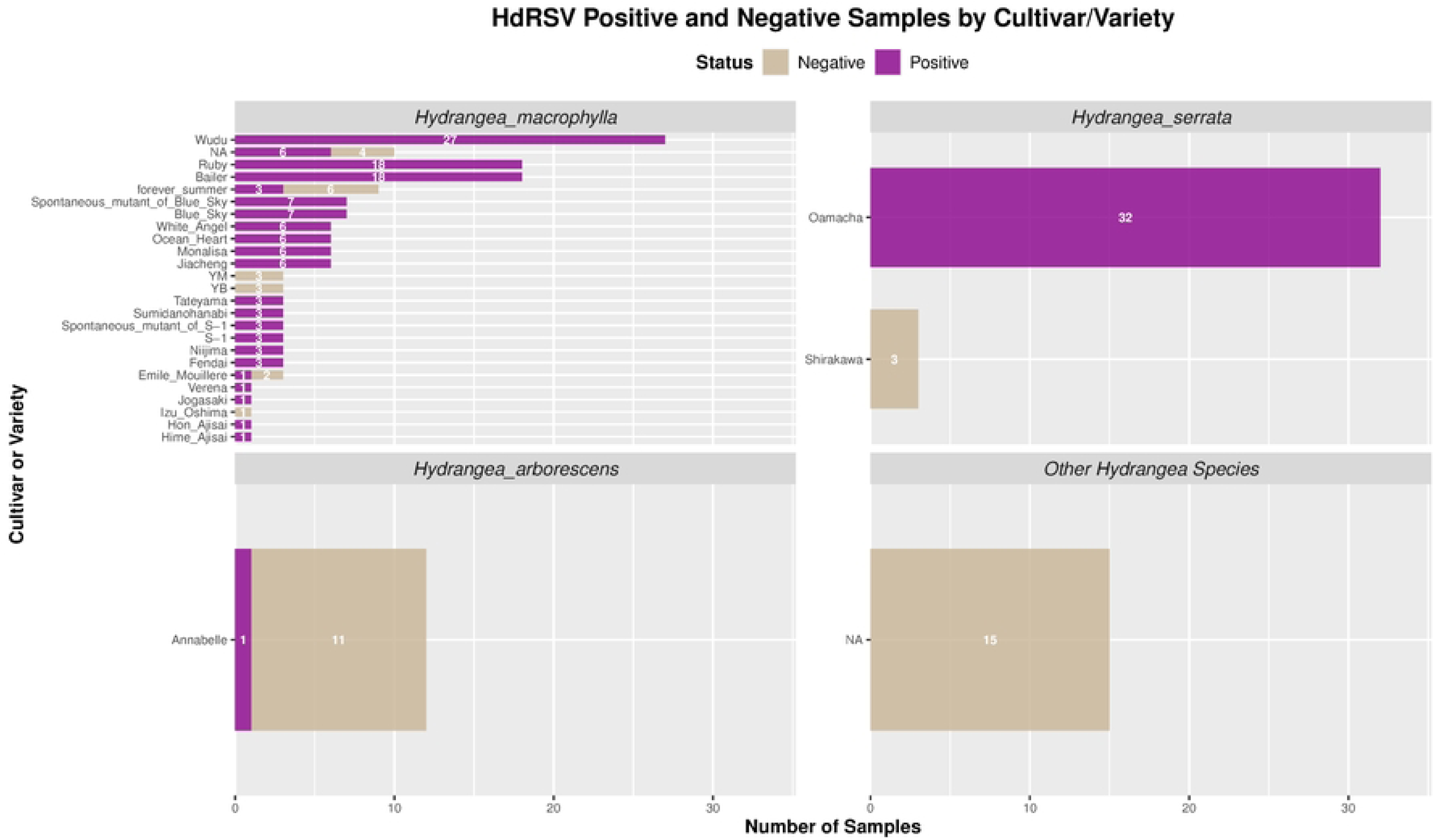
Prevalence of HdRSV across cultivars/varieties. Based on data from SRA and this study (including data from Colombia), this chart shows virus detection results for different cultivars, grouped by host species.

Lineage distribution varied by host species. HdRSV-L1 was the predominant lineage in *H. macrophylla* and the only one detected in *H. serrata*. Within our dataset, HdRSV-L2 was found exclusively in *H. macrophylla*. Remarkably, a single cultivar, Bailer, represented the only case in which both HdRSV lineages co-occurred. Sequences with incomplete replicase CDS coverage (<70%) were excluded from lineage assignment and are denoted as “NA” (Fig 8).

**Figure 8.**
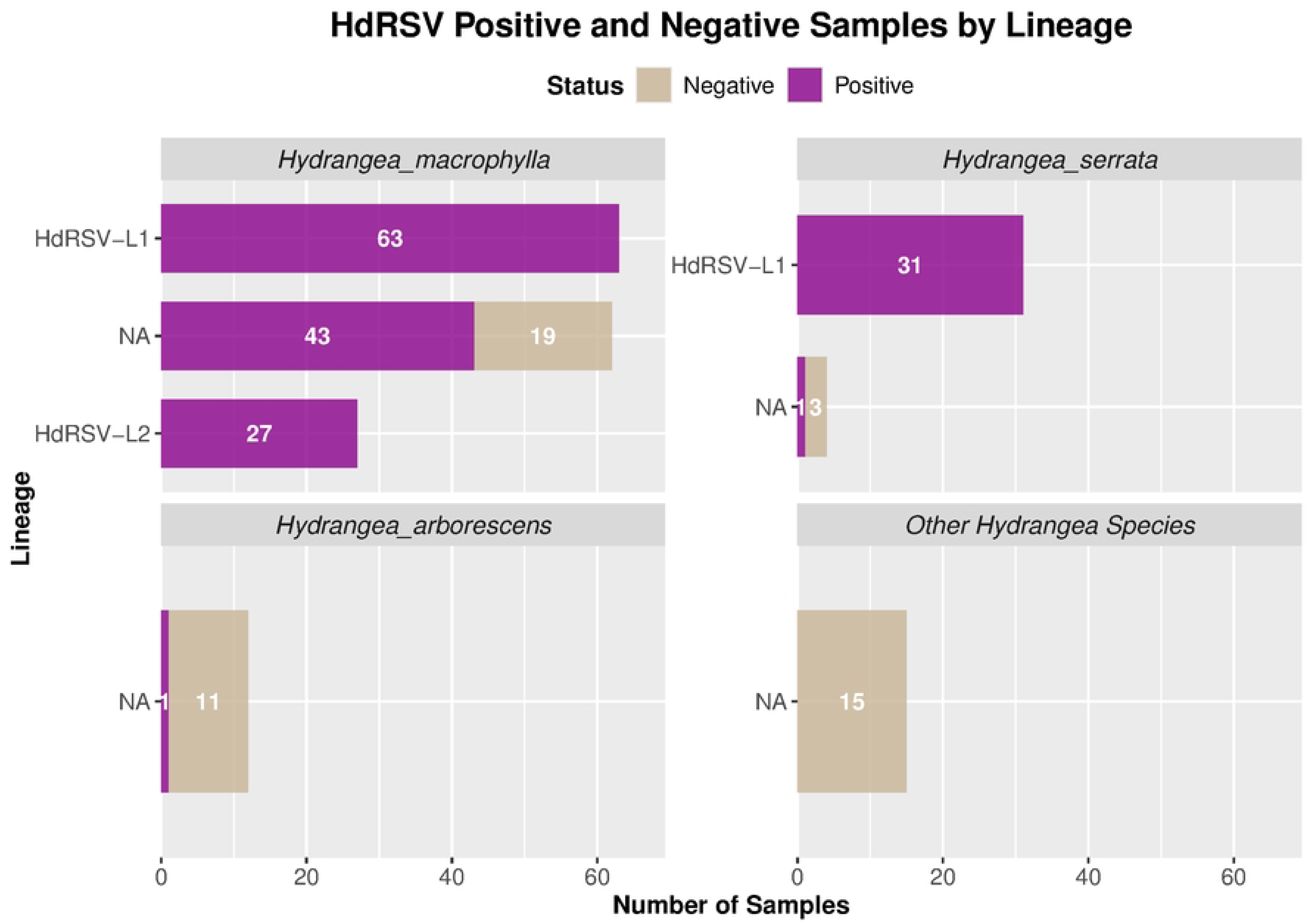
Distribution of HdRSV lineages across host species. Based on data from SRA and this study (including data from Colombia), this chart shows the classification of virus sequences into Lineage 1 (HdRSV-L1) and Lineage 2 (HdRSV-L2). Sequences with a replicase CDS coverage below 70% were excluded from lineage assignment and are categorized as ‘NA’ (Not Assigned).

### 3.7. HdRSV genome expression analysis

To compare the relative abundance of HdRSV across biological variables, viral genome expression levels were quantified. This analysis was restricted to *H. macrophylla* and *H. serrata*, for which a sufficient number of samples were available to support robust statistical inference.

Viral RNA abundance varied significantly across tissue types (Fig 9). The median viral load in root tissue was markedly higher than in all other tissues (bud, leaf, floral structures, and stem; all *p* < 0.001). This trend was most pronounced in *H. serrata*, where median root TPM values were approximately threefold higher than those observed in *H. macrophylla*. In *H. macrophylla*, the second highest viral load was detected in stem tissue, with a median TPM of 1,257. By contrast, floral structures, buds, and leaves exhibited substantially lower viral loads, with median TPM values below 51. In *H. serrata*, however, leaf tissues displayed notably elevated viral loads, reaching a median TPM of 24,541.

**Figure 9.**
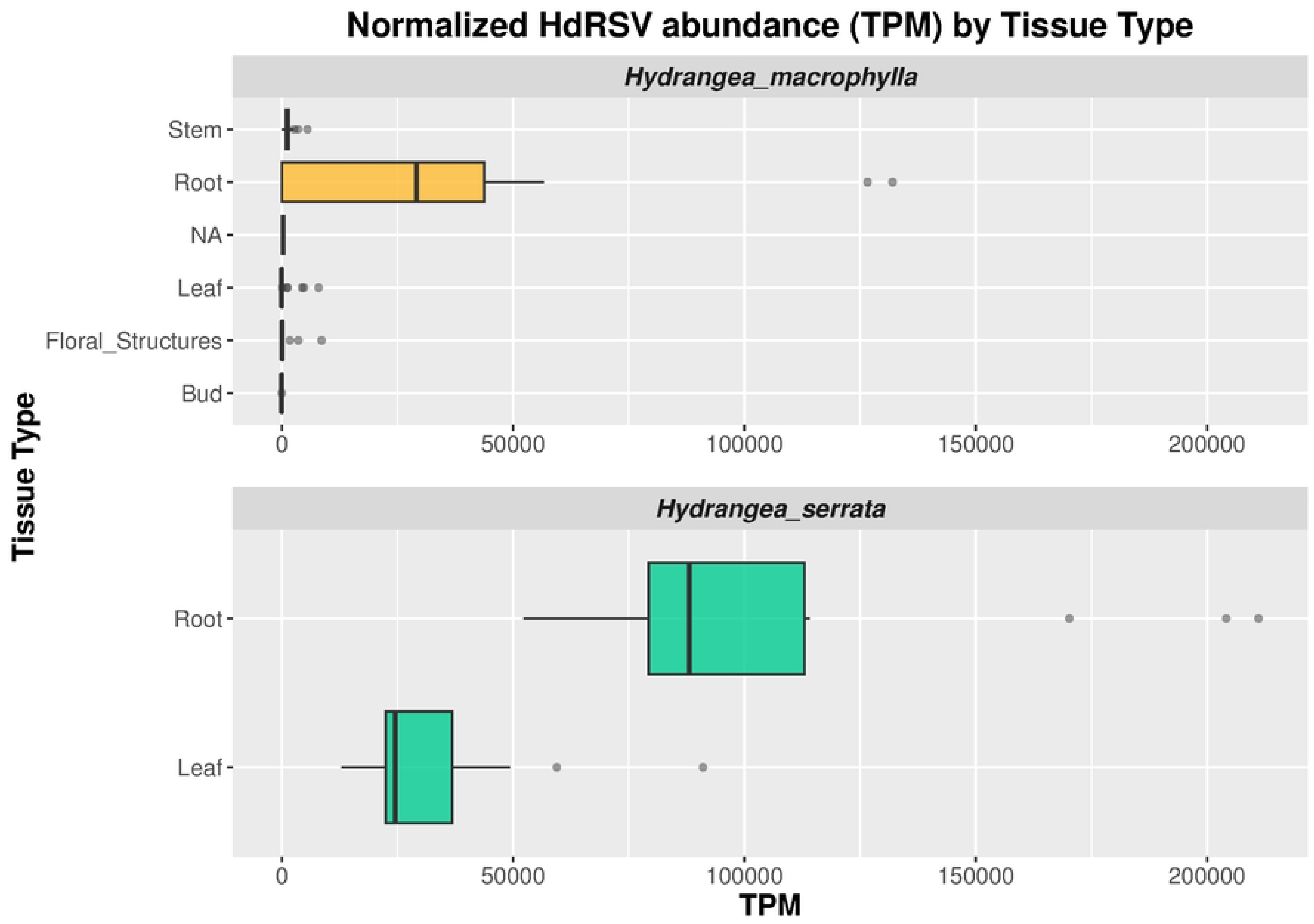
Expression levels of HdRSV across tissue types in *H. macrophylla* and *H. serrata*. This box-plot graph portrays the distribution of expression levels for HdRSV across different tissue types within the host species H. macrophylla and H. serrata. On the x-axis, expression values are presented in TPM (Transcripts Per Million). Each boxplot visualizes the median, interquartile range, and whiskers that delineate the data range.

Significant variation in viral abundance was also observed among cultivars (Fig 10). Oamacha, the only *H. serrata* cultivar included in the analysis, exhibited a significantly higher HdRSV load compared to multiple *H. macrophylla* cultivars. Within *H. macrophylla*, the cultivars Monalisa and Jiacheng showed significantly higher median TPM values than Bailer (Monalisa: p = 0.000358; Jiacheng: p = 0.00612).

**Figure 10.**
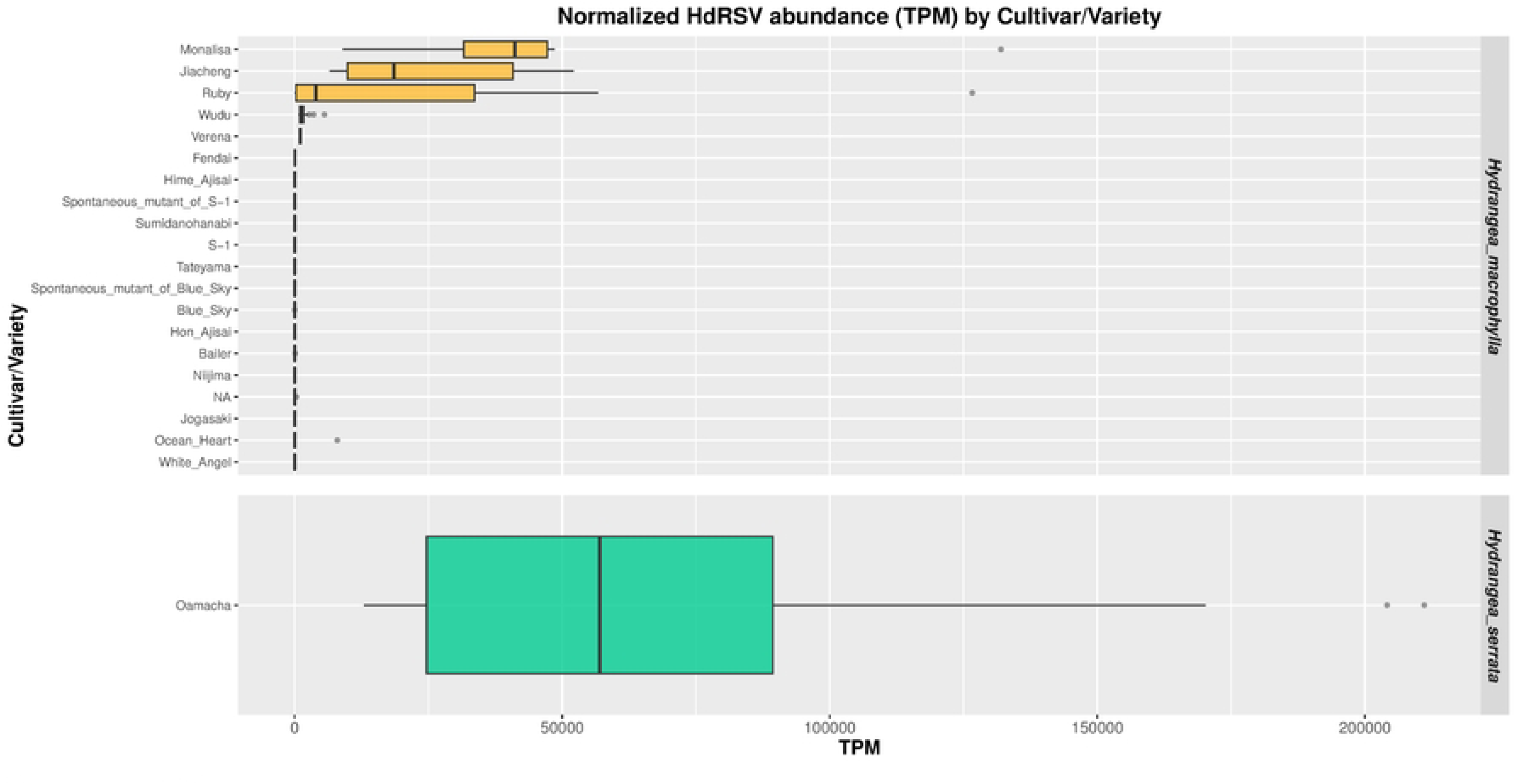
Expression levels of HdRSV across cultivars/varieties in *H. macrophylla* and *H. serrata*. This box-plot graph portrays the distribution of expression levels for HdRSV across different cultivars/varieties within the host species. On the x-axis, expression values are presented in TPM (Transcripts Per Million). Each boxplot visualizes the median, interquartile range, and whiskers that delineate the data range.

A direct comparison revealed that the overall median viral load was significantly higher in *H. serrata* than in *H. macrophylla* (p = 6.614e-15, Wilcoxon rank sum test with continuity correction). Furthermore, comparing HdRSV-L1 and HdRSV-L2, relative abundance, HdRSV-L1 was associated with higher viral loads, nearly more than 10 times (L1 median 2688 vs L2 218) (Wilcoxon test p-value: 0.02526662) (Fig 11).

**Figure 11.**
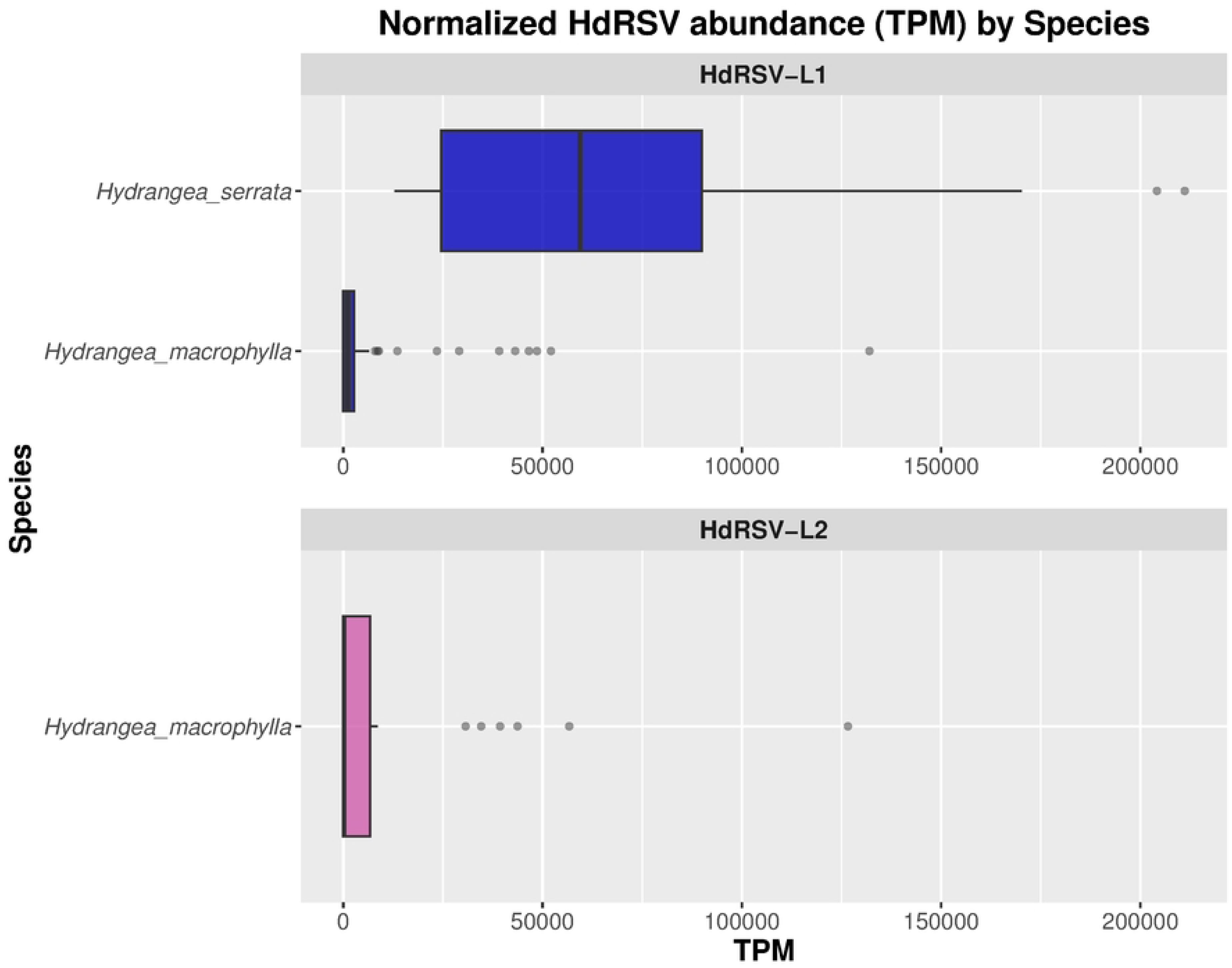
Expression levels of HdRSV across host species in lineages L1 and L2. This box-plot graph portrays the distribution of expression levels for HdRSV across the host species *H. macrophylla* and *H. serrata*, grouped by the viral lineages HdRSV-**L1** and HdRSV-**L2**. On the x-axis, expression values are presented in TPM (Transcripts Per Million). Each boxplot visualizes the median, interquartile range, and whiskers that delineate the data range.

The tissue-specific pattern of lineage abundance was consistent with the overall viral load distribution (Fig 12). Roots exhibited the highest TPM values for both lineages, with HdRSV-L1 showing the greatest accumulation, reaching a median value exceeding 52,000 TPM; however, the difference between lineages in this tissue was not significant. Leaves displayed the next highest viral abundance, again dominated by HdRSV-L1 (p = 0.0026).

**Figure 12.**
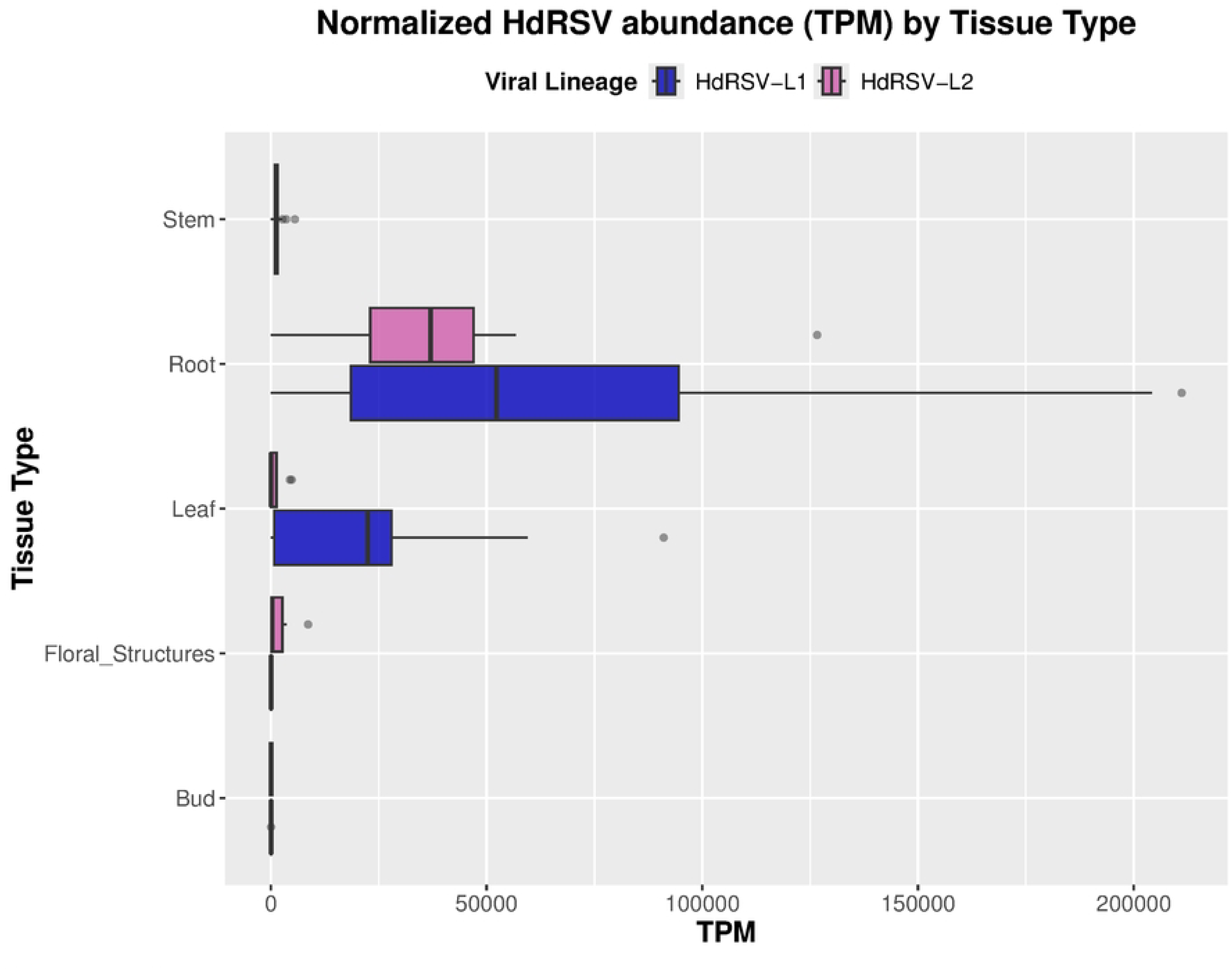
Expression levels of HdRSV-L1 and HdRSV-L2 across tissue types. This box-plot graph portrays the distribution of expression levels for HdRSV lineages L1 and L2 across different tissue types. On the x-axis, expression values are presented in TPM (Transcripts Per Million). Each boxplot visualizes the median, interquartile range, and whiskers that delineate the data range.

In contrast, floral structures and buds showed higher median TPM values for HdRSV-L2, although these differences were not statistically significant.

Analysis of lineage abundance by cultivar revealed that HdRSV-L1 reached its highest levels in Oamacha (TPM = 59,449), Monalisa (TPM = 41,142), and Jiacheng (TPM = 18,552), whereas HdRSV-L2 was most abundant in Ruby, although with a substantially lower TPM value (3,990) (Fig 13). Interestingly, both lineages were detected in biological replicates of the Bailer cultivar, representing the only instance of co-occurrence observed in the dataset.

**Figure 13.**
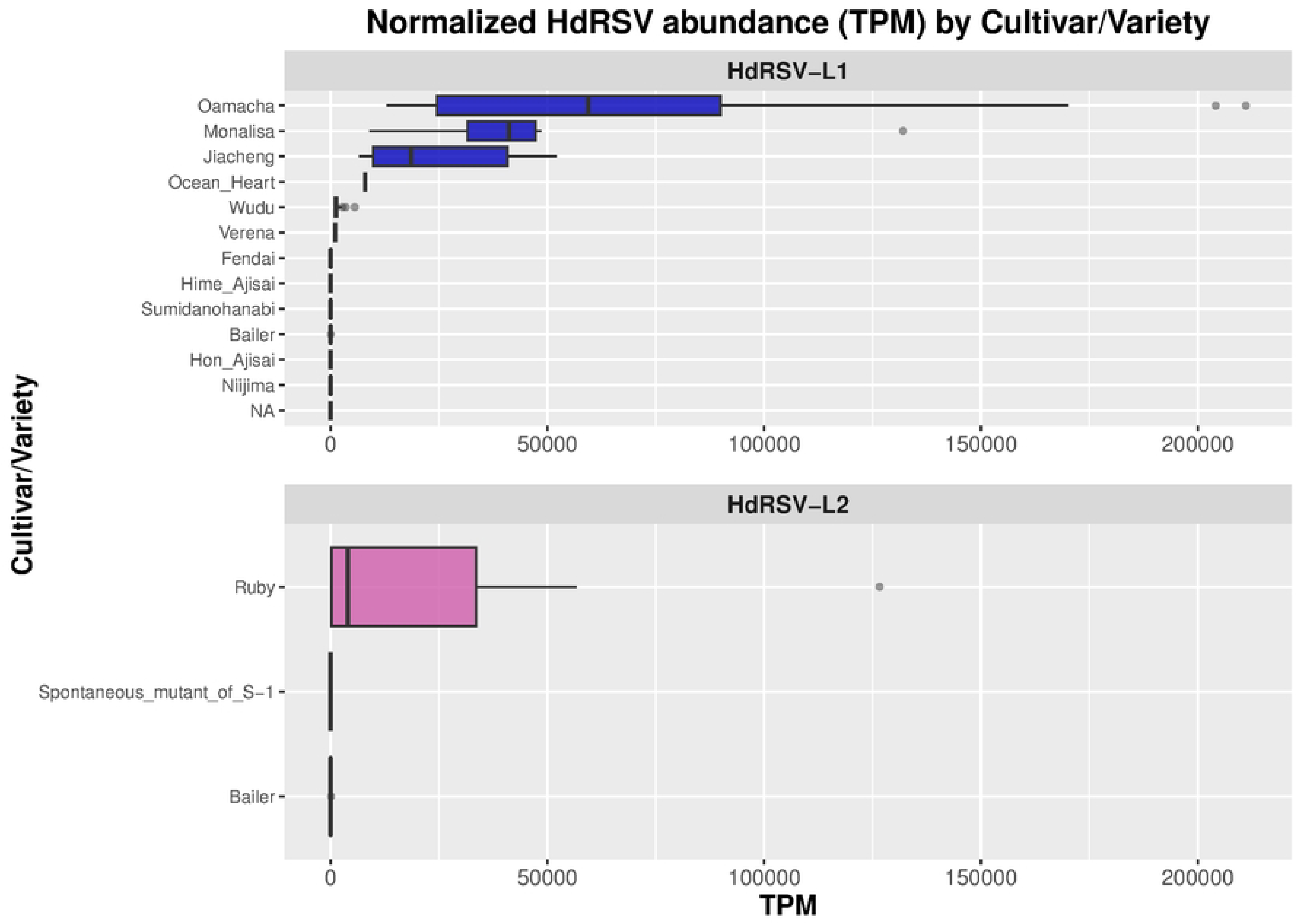
Expression levels of HdRSV across cultivars/varieties in lineages L1 and L2. This box-plot graph portrays the distribution of expression levels for HdRSV across different cultivars/varieties, grouped by the viral lineages HdRSV-L1 and HdRSV-L2. On the x-axis, expression values are presented in TPM (Transcripts Per Million). Each boxplot visualizes the median, interquartile range, and whiskers that delineate the data range.

### 3.8 HdRSV signature mutations

To investigate potential lineage-defining genetic differences in HdRSV, we analyzed the CDS of the viral replicase gene to identify fixed mutations distinguishing the two HdRSV lineages. Comparative analysis revealed several synonymous substitutions that, according to Treetime program, represent lineage-associated mutations. Although most were consistently specific to a given lineage, a few appeared reverted to the ancestral base in a minor fraction of the analyzed genomes. Remarkably, only a single non-synonymous substitution was identified in the replicase CDS, clearly segregating the two lineages. This substitution corresponds to a nucleotide change at position C1578T (replicase CDS coordinate based on reference NC_006943; second position of the codon), resulting in an amino acid replacement from threonine (ACC) to isoleucine (ATC). This mutation was present in approximately 90% of the genomes classified as lineage L2. Biochemically, this substitution represents a polar-to-nonpolar amino acid change: threonine (Thr) is a small, polar, uncharged residue containing a hydroxyl (–OH) group capable of hydrogen bonding and post-translational modification, whereas isoleucine (Ile) is a hydrophobic, branched-chain amino acid typically buried within protein cores or lipid-associated regions, contributing to hydrophobic packing and structural stability (Fig 14).

**Figure 14.**
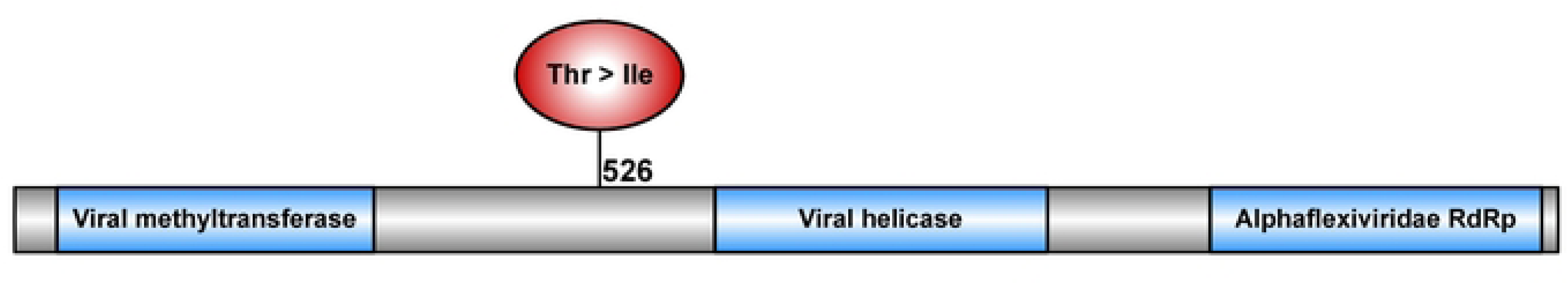
Schematic representation of the HdRSV replicase showing functional domains and the lineage-defining substitution Thr526Ile (C1578T), present in ∼90% of lineage L2 genomes. Figure generated with IBS (Liu et al., 2015).

## DISCUSSION

Previous studies on HdRSV were largely descriptive, focusing on isolated detections (Gadiou et al., 2010b; Hughes et al., 2005b; Akira Yusa et al., 2016). Only a handful of individual genome sequences have been reported globally. While such work helped assume its global presence and elucidated its genomic structure, it offered little information into population diversity or evolutionary dynamics. The scarcity of genomic data is particularly evident in the Americas, where all detections have been limited to short diagnostic fragments (Dória et al., 2011), leaving the virus’s evolutionary relationships unresolved. Our study shifts this paradigm through a large-scale genomic analysis. By leveraging high-throughput sequencing and public transcriptome data, we recovered 23 novel HdRSV genomes and generated the first genomic sequence from the Americas, a Colombian isolate, representing the most extensive genomic dataset currently studied for this virus. This effort dramatically expands the genetic documentation for HdRSV.

This expanded genomic resource enabled the first high-resolution analysis of HdRSV and supports a substantial refinement of the virus’s population structure. Our work addresses longstanding knowledge gaps by applying a combination of newly generated data from an infected *Hydrangea* plant in Colombia and an extensive survey of publicly available transcriptomes. Using Next-Generation Sequencing (NGS), we investigated HdRSV presence, evolutionary relationships, and patterns of viral expression. The breadth of sampling, including multiple cultivars, diverse geographic origins, and a range of host tissues, provided the depth and resolution necessary to characterize HdRSV more comprehensively and to establish a robust dataset for population-level analyses.

The phylogenetic analyses presented here consistently resolved two distinct and well-supported evolutionary lineages, designated HdRSV-L1 and HdRSV-L2. The strong statistical support recovered across the two reconstruction strategies indicates that this bipartite structure represents a stable feature of the HdRSV population in *H. macrophylla*. While different viral lineages have been reported within the genus *Potexvirus* (Fuentes et al., 2021; Wang et al., 2017), this study provides the first report of two distinct phylogenetic lineages for the species Hydrangea ringspot virus. This finding deepens our understanding of HdRSV’s genetic complexity and evolutionary history.

Geographical analysis revealed that both HdRSV lineages are widely distributed, with clear evidence of co-circulation in Germany and China. Notably, the detection of both lineages in multiple replicates of the same cultivar (Bailer) indicates that a single variety can be simultaneously susceptible to both viral strains, potentially facilitating their local maintenance and spread. However, the current dataset is heavily skewed toward Europe and East Asia, with minimal representation from the Americas and Africa. These sampling gaps limit our ability to draw definitive conclusions about global lineage distributions or potential region-specific adaptations. This underscores the urgent need for expanded viral surveillance in underrepresented regions, particularly given the phytosanitary risks inherent to the international ornamental plant trade (Groth-Helms and Zhang, 2024).

Plant viruses infect host tissues either through mechanical wounds or via biological vectors that deliver viral particles directly into cells, trespassing cell wall (Núñez-Muñoz et al., 2025). In the case of HdRSV, to date, no vector has been specifically associated with the transmission of this virus, and experimental evidence suggests that the virus is transmitted primarily through mechanical inoculation (Koenig, 1973). This mode of transmission aligns with our findings: a high infection rate (around 78%) and the observation that plants within the same cultivar are either all infected or all virus-free, cultivars with mixed status were extremely less common. Consequently, our results suggest that mechanical damage associated with horticultural handling, as previously reported, could play a predominant role in HdRSV transmission among cultivated hydrangeas (Groth-Helms and Zhang, 2024; Koenig, 1973).

The absence of HdRSV sequences in some samples within our evaluated transcriptomes suggests a contribution from both biological and technical factors. Biologically, these plants may be genuinely virus-free, possibly having been derived from uninfected propagation material. Alternatively, they could possess low-titer or incipient infections where a minimal viral replication rate fails to generate a discernible signal, making the few viral transcripts difficult to detect against the overwhelming predominance of host plant transcripts.

Viral tropism refers to the ability of a virus to infect and replicate within specific cell types, organelles, tissues, or hosts while encountering restrictions in others (Núñez-Muñoz et al., 2025). It has been described that actively replicating tissues, such as meristems and roots, can support higher viral replication rates. Conversely, mature leaf tissues provide a less favorable environment for viral multiplication due to their lower cellular replication activity and stronger innate immune responses, including a greater capacity to accumulate salicylic acid in response to infection (Ali et al., 2024; David et al., 2019; Pál et al., 2020).

Various host innate immune antiviral strategies are known to mediate viral tropism at different biological levels (Hipper et al., 2013; Mandadi and Scholthof, 2013). Our results revealed significantly higher HdRSV RNA loads in *H. serrata* compared to *H. macrophylla*, suggesting intrinsic differences in host–virus interactions. *H. serrata* may offer a more permissive cellular environment or possess less effective antiviral defenses. However, because all *H. serrata* sequences derived from a single cultivar, part of this difference could reflect cultivar-specific or environmental effects rather than species-wide susceptibility.

Plant viruses can colonize a broad range of host tissues (pantropism) or remain restricted to specific ones (Núñez-Muñoz et al., 2025). Our study suggests that HdRSV may have a pantropic infection pattern, being detected across all sampled tissues. However, viral accumulation was highest in roots, despite the virus not being soil-borne. This finding suggests that roots provide a particularly favorable environment for HdRSV replication or persistence, possibly due to tissue-specific physiological conditions. Although it was established that HdRSV enter plant vascular tissues through mechanical inoculation, the pronounced viral load in roots indicates that additional mechanisms might facilitate its movement or accumulation in underground tissues. Previous studies have reported that at least 39 plant viruses, including members of *Alphaflexiviridae* (which includes the genus *Potexvirus*), *Benyviridae*, *Bromoviridae*, *Closteroviridae*, *Potyviridae*, *Secoviridae*, *Solemoviridae*, *Tombusviridae*, and *Virgaviridae*, are capable of invading xylem tissues (Núñez-Muñoz et al., 2025; Sun et al., 2022). Despite this ability, their accumulation patterns often differ, and their movement can occur independently of phloem sap flow. Current models suggest that these viruses may initially infect cells adjacent to immature xylem, subsequently entering the transpiration stream and disseminating through guttation. However, how viruses move from mature xylem vessels into living plant tissues remains poorly understood (Núñez-Muñoz et al., 2025; Sun et al., 2022). In the case of HdRSV, the high viral RNA levels detected in roots, despite no confirmed evidence of a soil vector or direct root inoculation, expands the conventional understanding of virus spread and tissue tropism in *Potexvirus* species.

Visible root symptoms are rare, even in soil-borne viral infections (Núñez-Muñoz et al., 2025), which explains the limited information available about root-specific infections, particularly in non–soil-borne viruses. Taken together, our findings suggest that HdRSV exhibits a distinctive pattern of tissue tropism characterized by pantropism with a high root accumulation, potentially driven by both host-specific and physiological factors that remain to be elucidated.

At the lineage level, the significantly higher overall abundance of HdRSV-L1, particularly in *H. serrata*, suggests this lineage may have a replication or transmission advantage over HdRSV-L2. Furthermore, the detection of both HdRSV-L1 and HdRSV-L2 in the ‘Bailer’ cultivar provides evidence of mixed HdRSV infections in hydrangeas. This co-occurrence might create an opportunity for genetic recombination between lineages, a known mechanism for the emergence of novel viral variants with altered virulence. Confirming co-infection within individual plants and assessing the potential for recombination should be a priority for future research.

Altogether, these findings—encompassing large-scale genomic recovery, well-supported phylogenetic structure, host-associated lineage patterns, and tissue-specific differences in viral accumulation—provide an expanded framework for understanding the evolution, epidemiology, and persistence of HdRSV in ornamental plant systems

---

**Supplementary Figure 1.**
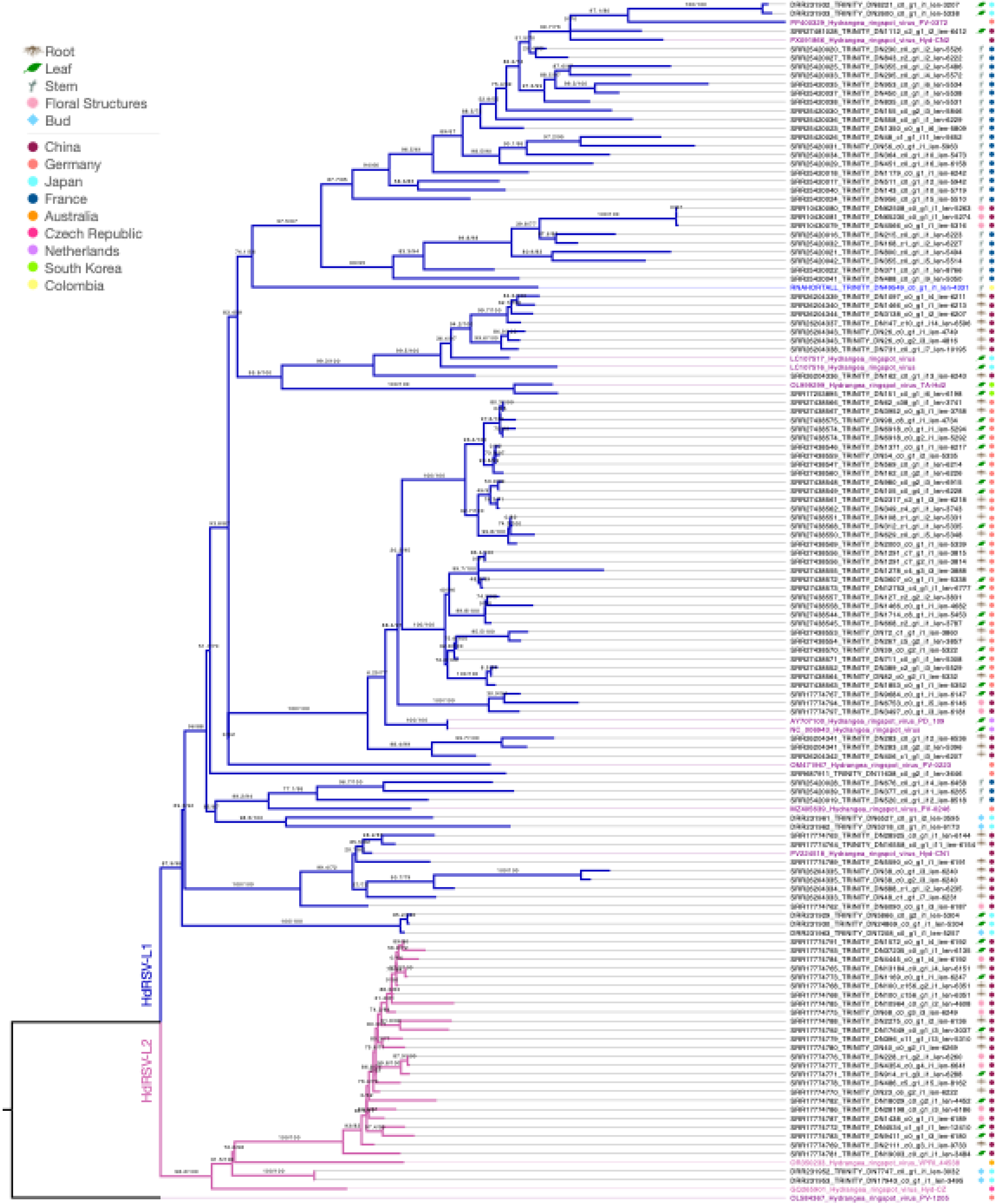
Phylogenetic analysis of HdRSV variants. The unrooted maximum-likelihood tree was constructed using codon-based models and the replicase coding sequence (CDS) from HdRSV isolates with ≥70% coverage, together with 13 reference genomes from GenBank. Branch lengths are proportional to genetic distance. Support values at nodes represent ultrafast bootstrap (UFB, 5000 pseudoreplicates) and approximate likelihood ratio test (aLRT) values.

**Supplementary Table 1** Descriptions of the SRA accessions used in this study, including HdRSV (Hydrangea ringspot virus, *Potexvirus hydrangeae*) mapping results, normalized TPM counts, and associated geographic and cultivar metadata.

